# Rhodopsin-based voltage imaging tools for use in excitable cells of *Caenorhabditis elegans*

**DOI:** 10.1101/544429

**Authors:** Negin Azimi Hashemi, Amelie C. F. Bergs, Rebecca Scheiwe, Wagner Steuer Costa, Jana F. Liewald, Alexander Gottschalk

## Abstract

Genetically encoded voltage indicators (GEVIs) based on microbial rhodopsins utilize the voltage-sensitive fluorescence of the all-trans retinal (ATR) cofactor, while in electrochromic (eFRET) sensors, donor fluorescence drops when the rhodopsin acts as depolarization-sensitive acceptor. We systematically assessed Arch(D95N), Archon, and QuasAr, as well as the eFRET sensors MacQ-mCitrine and QuasAr-mOrange, in *C. elegans*. ATR-bearing rhodopsins reported on voltage changes in body wall muscles (BWMs) and the pharynx, the feeding organ, where Arch(D95N) showed ca. 125 % ΔF/F increase per 100 mV. The ATR fluorescence is very dim, however, using the retinal analog dimethylaminoretinal (DMAR), it was boosted 250-fold. eFRET sensors provided sensitivities of 45 % to 78 % ΔF/F per 100 mV, induced by BWM action potentials (APs). All sensors reported differences in muscle depolarization induced by a voltage-gated Ca^2+^-channel mutant. Optogenetically evoked de-or hyperpolarization of motor neurons increased or eliminated AP activity and caused a rise or drop in BWM sensor fluorescence. Last, we could analyze voltage dynamics across the entire pharynx, showing uniform depolarization but compartmentalized repolarization of anterior and posterior parts. Our work establishes all-optical, non-invasive electrophysiology in intact *C. elegans*.

## INTRODUCTION

Activity of excitable cells like muscles and neurons can be measured by electrophysiology, Ca^2+^-or voltage imaging^1-7^. While electrical measurements provide the highest sensitivity and temporal accuracy, imaging methods are much more versatile for applications in living animals and for recording multiple cells simultaneously. Genetically encoded Ca^2+^ indicators (GECIs) nowadays cover most of the visible spectrum with comparably narrow spectral width, thus enabling multiplexing with other optical tools^8-10^. Furthermore, GECIs were improved immensely since their first reporting in the late 1990’s, to provide several thousand-fold of fluorescence increases upon Ca^2+^-binding^5^. With differential kinetic properties, some GECIs enable detection of single action potentials (APs). They are thus widely used in neuroscience, yet in many neuron types Ca^2+^ imaging does not reflect spiking dynamics, and does not permit to assess high frequency APs or sub-threshold voltage fluctuations^11^. Also, Ca^2+^ is an indirect measure of neuronal activity, and neuronal depolarization is not always accompanied by a Ca^2+^ increase^12^. Finally, GECIs are less-well suited to detect a hyperpolarization of cells, as there is usually no decrease of Ca^2+^ below the basal cytosolic concentration. Thus, voltage imaging tools (genetically encoded voltage indicators - GEVIs) are an important addition to the toolbox of sensors for excitable cell activity^11,13-18^.

The development of GEVIs started later than that of the GECIs, and several different designs were explored that couple detection of voltage changes to fluorescence changes. These range from FRET sensors to circularly permuted GFP, similar as for GECIs^4,5^. Yet, such tools for a long time reached only a few percent in fluorescence change per 100 mV of membrane voltage change, thus making detection of single APs a challenge. More recently, microbial rhodopsins were found to exhibit a voltage-dependent fluorescence of their chromophore, retinal^17,19^. The fluorescence signal change was higher than for the other protein-based GEVIs (ca. 20-40 % per 100 mV), and could be improved by several mutagenesis screens^13,18^. However, the absolute fluorescence signals of the rhodopsins are very small. In particular, the voltage sensitivity of the fluorescence, due to the nature of the process requiring more than one photon to be absorbed^20^, becomes appreciable only at very high light intensities. Thus, electrochromic Förster resonance energy transfer (eFRET) sensors were developed that couple fluorescence of a normal fluorescent protein (FP) to the rhodopsin. The latter is acting as a FRET acceptor upon depolarization, thus quenching the (much stronger) fluorescence of the FP, which acts as a FRET donor^14,16^. Several variants of eFRET sensors have been reported, which are combinations of variants of archaerhodopsin (QuasAr^18^, Archon^13^) or other proton pumps like Mac (from *Leptoshaeria maculans*) or Ace2N, derived from the *Acetabularia* rhodopsin proton pump^14,21,22^. Each of these proteins is coupled with specific linkers to fluorescent proteins like mOrange, mCitrine, mNeon, or mRuby3, depending on the absorption spectrum of the respective rhodopsin, to achieve maximal FRET efficiency, and to achieve imaging in different spectral regions.

An additional way to improve voltage-dependent fluorescence of the rhodopsin may be to alter the chemical properties of the retinal chromophore. We previously used retinal analogs to reconstitute function of microbial rhodopsin optogenetic tools, and to alter their characteristics^23^. During these studies, we observed that some retinal analogs conferred a much higher fluorescence to the rhodopsin. Here, we surveyed a range of GEVIs based on microbial rhodopsins for use in excitable cells of the nematode *Caenorhabditis elegans*, an important model system in physiology, and in molecular, cellular and behavioral neurobiology. We demonstrate that eFRET sensors are robust tools allowing to analyze voltage signals with little experimental effort, and that the retinal analog <u>dim</u>ethyl<u>a</u>minoretinal (DMAR) strongly improves absolute fluorescence signals, however, at the cost of lower fluorescence changes. We show that direct voltage-dependent retinal fluorescence can be measured in the infrared at high excitation intensity, and that membrane voltage can be set concomitantly with rhodopsin tools for depolarization or hyperpolarization. Alterations in AP amplitude and duration could be robustly detected in gain-of-function mutants affecting the L-type Ca^2+^ channel EGL-19 in a spontaneously active muscular pump, the pharynx, allowing also correlation with the timing of pump events. Last, GEVIs enabled to visualize the spatiotemporal compartmentalization of voltage changes in this muscular structure.

## RESULTS

### Archaerhodopsin Arch(D95N) equipped with ATR shows dim, fluctuating fluorescence in muscle, that is enhanced by DMAR

To explore the potential of analyzing muscle voltage changes via rhodopsin/retinal fluorescence, we expressed Arch(D95N)^17^, QuasAr (an improved mutated variant of Arch^18^) and Archon^13^, a further evolved version in body wall muscle (BWM) cells, or in pharyngeal muscle, and supplemented the animals with all-*trans* retinal (ATR). Under an epi-fluorescence microscope equipped with a 100W HBO lamp for excitation and a sCMOS camera, very dim fluorescence could be observed in the range of 700 nm. Using a 637 nm laser, and an EMCCD camera, fluorescence was more readily observable (**Fig. 1A, B**).

**Figure 1.**
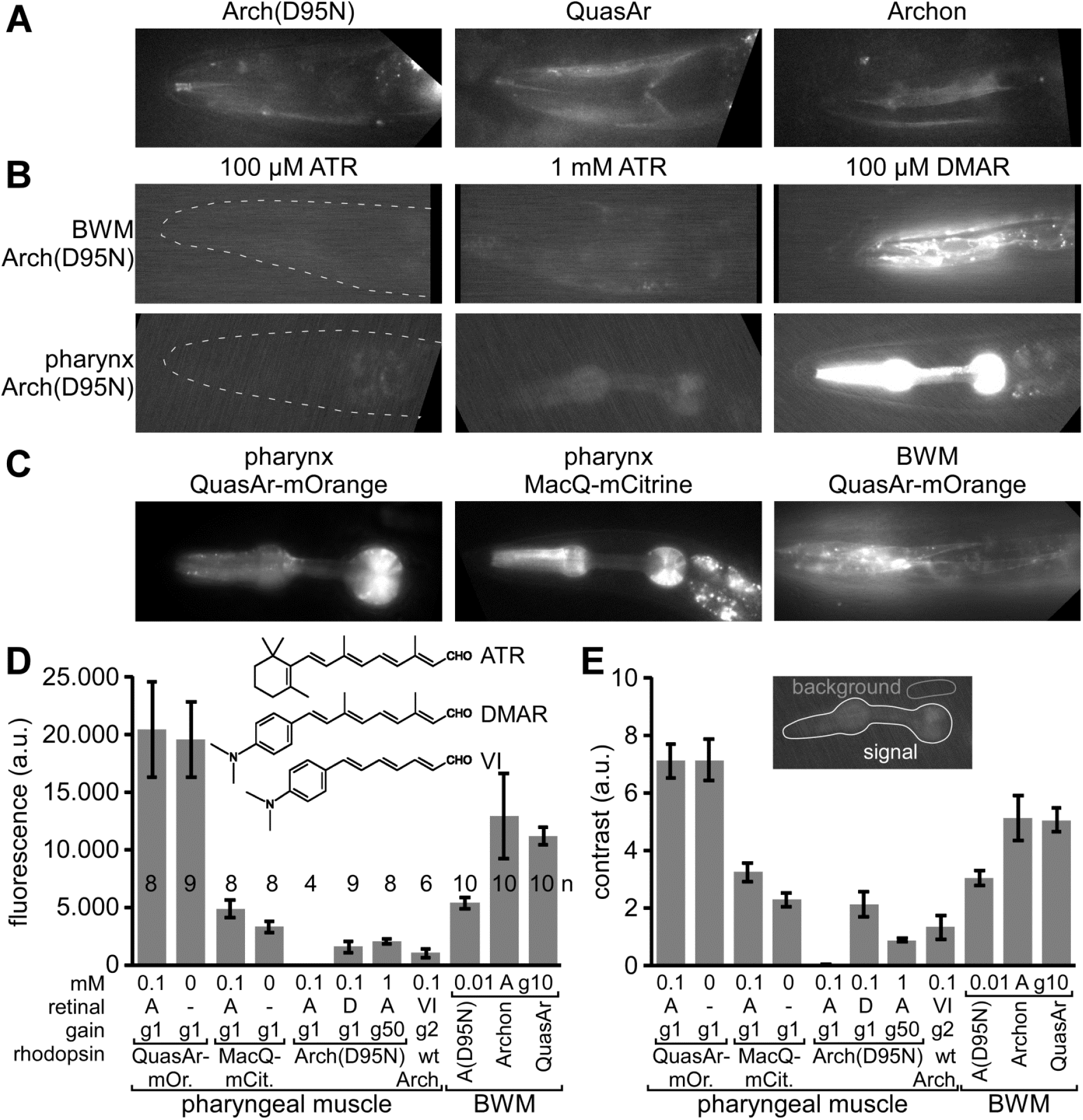
Expression of rhodopsin voltage sensors in *C. elegans* muscle cells: **A)** Expression and imaging of retinal fluorescence in body wall muscle cells in the head, of (from left to right) Arch(D95N), QuasAr, and Archon. Scale bar = 10 µm. **B)** Expression and imaging of Arch(D95N) in BWMs (top row) and in the pharyngeal muscle (bottom row), complemented with ATR (left and middle, two different concentrations) and with the retinal analog DMAR (right; see inset in (D) for chemical structures of ATR, DMAR and retinal analog VI). Dashed outline in left panels indicates worm head. **C)** Expression and imaging of eFRET voltage sensors MacQ-mCitrine and QuasAr-mOrange in pharyngeal muscle and in BWMs, as indicated. **D, E)** Characteristics of basal fluorescence for the sensors expressed in the pharynx or in BWMs, as indicated. Fluorescence intensity (D) as average grey values of a ROI enclosing the entire pharynx and contrast (E) as the ratio of signal over mean fluorescence of a control ROI were acquired (20 ms exposure time) with different gain (indicated as ‘g’ and a number). The rhodopsins were supplemented either with ATR or DMAR, or analog VI (‘A’, ‘D’, ‘VI’) of the indicated concentrations (in mM). Shown are mean±SEM. Number of animals imaged is indicated in (D). In A-C, anterior is to the left.

In comparison to Arch(D95N), the fluorescence of QuasAr and Archon was 2.4 and 2.1-fold more intense, respectively, and showed a 1.7-fold higher contrast over background fluorescence (in BWMs; **Fig. 1D, E**). In immobilized animals, this fluorescence showed intensity fluctuations in the range of 20-25 % ΔF/F that likely represent voltage changes (**Supp. Fig. 1**), possibly corresponding to APs as reported earlier for *C. elegans* BWM cells^24,25^.

**Supplementary Figure S1:**
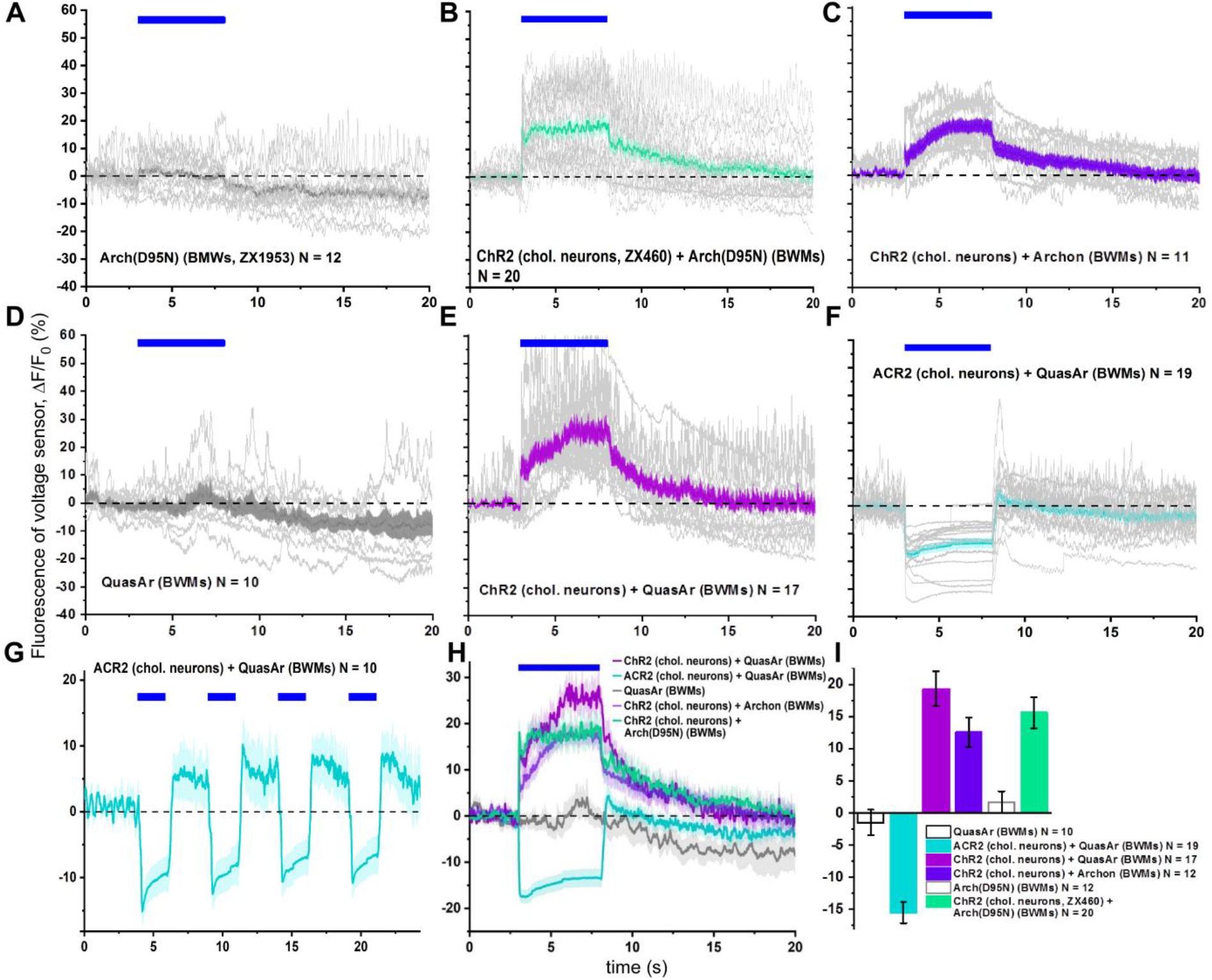
Summary of individual and mean voltage fluorescence traces for various rhodopsin voltage sensors expressed in BWMs. Related to Figs. 1 and 3. **A)** Arch(D95N)-ATR, spontaneous activity. **B)** Arch(D95N)-ATR with ChR2 expressed in cholinergic neurons. **C)** Archon with ChR2 expressed in cholinergic neurons. **D)** QuasAr, spontaneous activity. **E)** QuasAr with ChR2 expressed in cholinergic neurons. **F, G)** QuasAr with ACR2 expressed in cholinergic neurons. **H)** Summary of mean fluorescence changes in (B, C, E, F). Blue bars in (A-H) indicate photo-stimulation periods. **I)** Statistical analysis of the mean fluorescence changes of data shown in H, averaged across the 5 s stimulation periods.

When we supplemented the animals with the retinal analog DMAR^23^ (**Fig. 1D**, inset), we observed a ca. 250-fold higher absolute fluorescence (in pharyngeal muscle; **Fig. 1B, D**), and similarly, when we used retinal analog VI, fluorescence levels increased 81-fold (gain-corrected; note this was in Arch wt, in which this analog does not mediate function, and thus does not hyperpolarize the cell; analog VI could not be incorporated by Arch(D95N)). This was accompanied by higher signal contrast (138-fold and 44-fold higher for DMAR and analog VI, respectively; **Fig. 1E**). The fluorescence yield of Arch(D95N) could also be increased by using 10-fold higher ATR concentration during animal cultivation, and a 50x higher camera gain during imaging (**Fig. 1B, D**), resulting in fluorescence grey values that were similar to those observed for DMAR. Increased fluorescence due to DMAR or increased ATR and gain was also observed in BWM cells (**Fig. 1B**). Last, we also expressed eFRET sensors, QuasAr-mOrange and MacQ-mCitrine, in pharyngeal muscle or BWM (**Fig. 1C**). Since here, fluorescence of canonical FPs is observed, their absolute fluorescence was much higher than for any of the rhodopsin-only sensors (ca. 3000-fold and 800-fold higher fluorescence than Arch(D95N) equipped with ATR, for QuasAr-mOrange and MacQ-mCitrine, respectively; **Fig. 1D**), and likewise, their contrast level compared to background was much higher (460-and 211-fold higher, respectively; **Fig. 1E**).

### Intrinsic activities in the pharynx and neuromuscular systems are accompanied by fluorescence changes of rhodopsin GEVIs

To assess if the fluorescence of the sensors is also voltage-dependent, we observed it during intrinsic activity of the muscular systems. For the pharynx, we immobilized animals such that they would still exhibit pharyngeal pumping, when pre-treated with serotonin (to mimic the presence of food). As exemplified for QuasAr-mOrange, fluorescence changes are apparent upon pharyngeal muscle contraction as a drop in the fluorescence that, importantly, was only observed when the protein was supplemented with ATR, thus yielding a functional eFRET acceptor (**Fig. 2A, B, Supp. Video 1**; the fluorescence of the terminal bulb (TB; grinder region) was used to visualize pharyngeal contractions as the TB lumen opens up).

**Figure 2.**
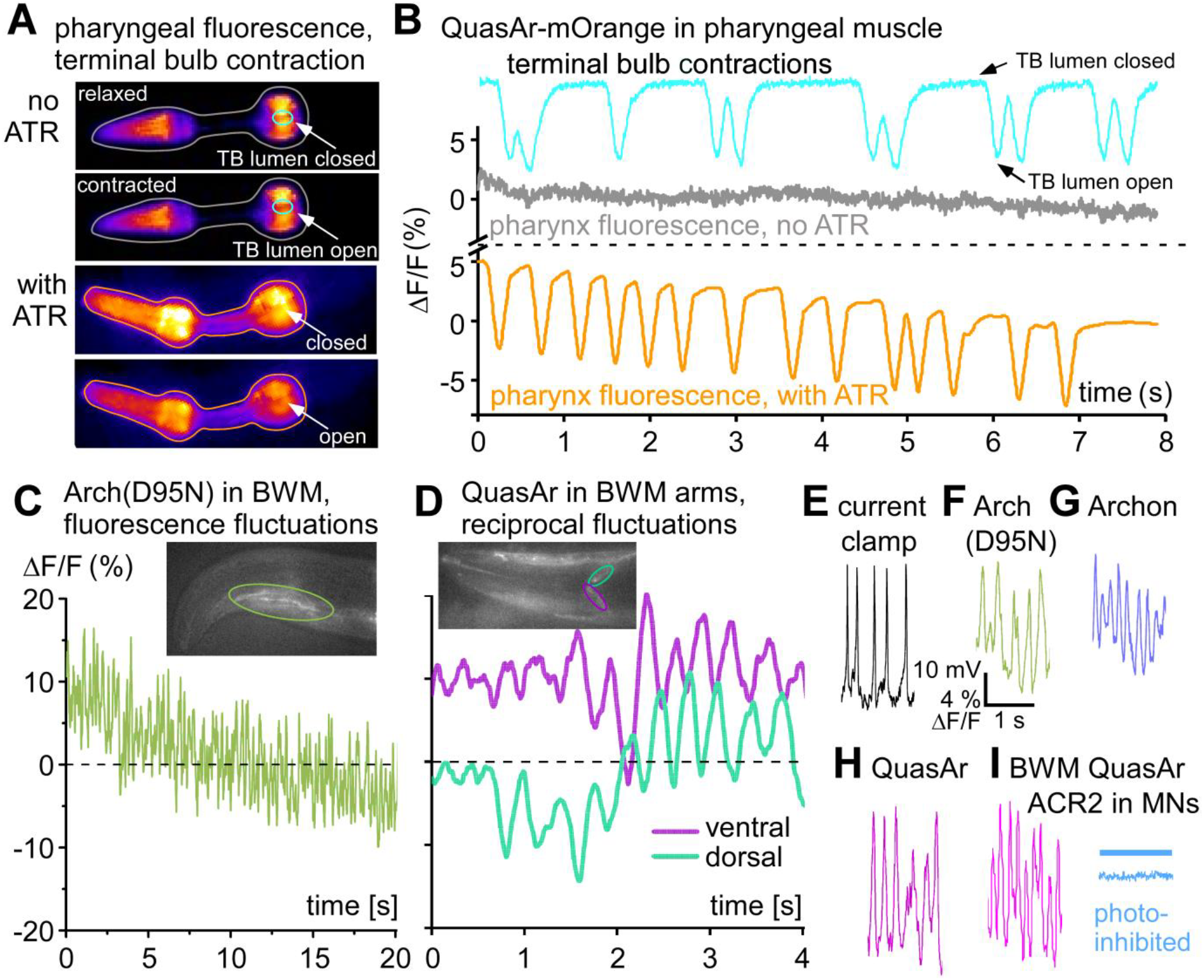
Rhodopsin voltage sensors detect intrinsic voltage changes in muscular tissues: **A)** Expression of the QuasAr-mOrange eFRET sensor in pharyngeal muscle, false-color representation of fluorescence intensity. Upper two images, animal raised without ATR; still images from a video depicting relaxed and contracted states. ROIs are drawn for the fluorescence of the entire organ (grey), as well as for the terminal bulb (TB) lumen in the grinder region (cyan; time series for both ROIs are shown in B). Lower two images are from a video of an animal that was supplemented with ATR (voltage fluorescence ROI is drawn in orange). A drop in fluorescence is apparent upon pharyngeal contraction, the lumen of the TB opens. **B)** ΔF/F fluorescence time traces (acquired at 189 fps, 1 ms exposure) of the ROIs indicated in (A), in %. The cyan graph shows fluctuations of the fluorescence signal due to opening and ‘darkening’ of the TB lumen, closed and open ‘states’ are indicated. Without ATR, no change in fluorescence is observed despite pumping (grey trace; from same video as for TB contraction). Instead, animal kept on ATR shows strong fluorescence drop upon pharyngeal activity. See also **Supp. Video 1**. **C)** Arch(D95N) fluorescence imaged in a BWM in the head of the animal (ROI indicated in inset, acquired at 158 fps, 2 ms exposure). 10-15 % fluctuations at high rate are observed, as well as signal beaching. **D)** QuasAr imaging in BWM, dorsal and ventral muscle arm in the nerve ring (ROIs indicated in turquoise and magenta, respectively), ΔF/F fluorescence shows dorso-ventral alternation, as expected for muscle activation during undulatory locomotion. See also **Supp. Video 2**. **E)** Action potentials recorded in dissected BWM cell, under current clamp, exhibiting typical amplitudes and frequency. **F-H)** Fluorescence fluctuations of similar frequencies in BWM cells expressing the rhodopsin sensors Arch(D95N) (F), Archon (G) and QuasAr (H); each acquired at 158 fps, 2 ms exposure time. See also **Supp. Videos 3, 4**. **I)** AP-like activity imaged in QuasAr expressing BWM cell, is inhibited by ACR2-mediated optogenetic inhibition of cholinergic motor neurons (MNs), upstream of the BWM (blue trace, blue bar indicates illumination).

**Supplementary Video 1:** QuasAr-mOrange in pharynx, spontaneous pumping.

For Arch(D95N) in BWM cells, we observed fluorescence fluctuations that occurred at a rapid time scale (**Fig. 2C**), reminiscent of BWM APs appearing spontaneously in dissected animals, that can be recorded by current clamp electrophysiology (**Fig. 2E**). To verify that the fluorescence fluctuations were in response to muscular depolarization, we analyzed the muscle arms of QuasAr expressing BWM cells in the head (**Fig. 2D; Supp. Video 2**). Here, we observed fluorescence fluctuations in a dorso-ventrally reciprocal fashion, as would be expected for the control of muscles during undulatory locomotion of the animal. As we and others showed previously, such reciprocal activity is apparent in immobilized animals^26^, even though at a slower time scale than expected for freely moving animals. We observed fluctuations of about 10-20 %ΔF/F for all the rhodopsin-based voltage sensors tested (Arch(D95N), Archon, QuasAr; **Fig. 2F, G, H; Supp. Fig. 1; Supp. Videos 3, 4**). Importantly, when we expressed and co-activated the natural anion-selective channelrhodopsin ACR2^27,28^ in cholinergic motor neurons (MNs), fluorescence fluctuations in BWM cells expressing QuasAr were absent (**Fig. 2I; Supp. Fig. 1D**), verifying that the fluctuations were evoked by presynaptic cholinergic MNs that depolarized BWM cells via acetylcholine (ACh) signaling.

**Supplementary Video 2:** QuasAr in muscle arms, dorso-ventrally alternating intrinsic activity.

**Supplementary Video 3:** QuasAr in BWMs, intrinsic activity.

**Supplementary Video 4:** QuasAr in BWMs, intrinsic activity.

### ChR2-mediated depolarization and ACR2-mediated hyperpolarization of MNs enables calibration of rhodopsin voltage signals in BWMs

We sought to estimate how voltage-induced fluorescence changes compare to electrical depolarization voltages. The APs of BWM cells can be evoked by stimulated release of ACh from MNs. When recorded by current clamp, ChR2 stimulation evoked mean APs of 29.15 mV (n=8; **Figs. 2E; 3A, B, I**). Given the enhanced imaging possibilities using laser excitation, we tested the possibility to achieve imaging in BWM cells concomitant with photo-depolarization and -hyperpolarization of pre-synaptic MNs. To this end, we expressed ChR2 or ACR2 in cholinergic MNs, as well as three different voltage sensors in BWMs (Arch(D95N), QuasAr, and Archon). Fluorescence for all three proteins was well visible along the BWM plasma membrane of the respective transgenic animals, following 637 nm laser excitation (**Fig. 1A**). Imaging basal fluorescence signals (i.e. without actuator-mediated stimulation) revealed spontaneous brief increases of fluorescence of ≈15 % ΔF/F for QuasAr, and ≈10 % for Archon and for Arch(D95N) (**Fig. 2F-H; Supp. Fig. 1**). These lasted for ca. 50-100 ms, and were reminiscent of APs measured by electrophysiology in BWM cells (**Fig. 2E**). During prolonged recordings (25 s), QuasAr and Archon showed robust signals that showed little alteration in baseline, while Arch(D95N) exhibited photo bleaching (**Supp. Fig. 1A, D, H**).

When we used ChR2 stimulation of MNs, but not in animals lacking ChR2, mean evoked increased ΔF/F signals, corresponding to depolarization, were observed for all sensors: ≈22.7 % for QuasAr and Arch(D95N), and ≈13.3 % for Archon, for the first evoked peak (**Fig. 3C-F, I;** for Archon, see **Supp. Fig. 1C, H, I**). Interestingly, trains of APs could be observed that were overlaid on the overall increased fluorescence rise. When averaged over the 5 s stimulus period, signals were ≈21.4 % for QuasAr, ≈14.6 % for Archon, and ≈17% for Arch(D95N). When we used ACR2 for hyperpolarization of cholinergic MNs, the fluorescence of QuasAr dropped (repeatedly) by about 14.6 %, and no AP-like spikes, overlaid on the fluorescence trace, were observable (**Figs. 2I, 3G-I; Supp. Fig. 1F-I**). These measurements allow a rough calibration of the observed fluorescence voltage signals, namely ca. 78 % ΔF/F per 100 mV for QuasAr and Arch(D95N), ca. 46 % ΔF/F per 100 mV for Archon. This data is in a similar range as reported in the literature (ca. 45% ΔF/F per 100 mV for Arch(D95N)^17^, QuasAr^18^ and Archon^13^), though it appears as if the first two sensors perform better in *C. elegans* tissue.

**Figure 3.**
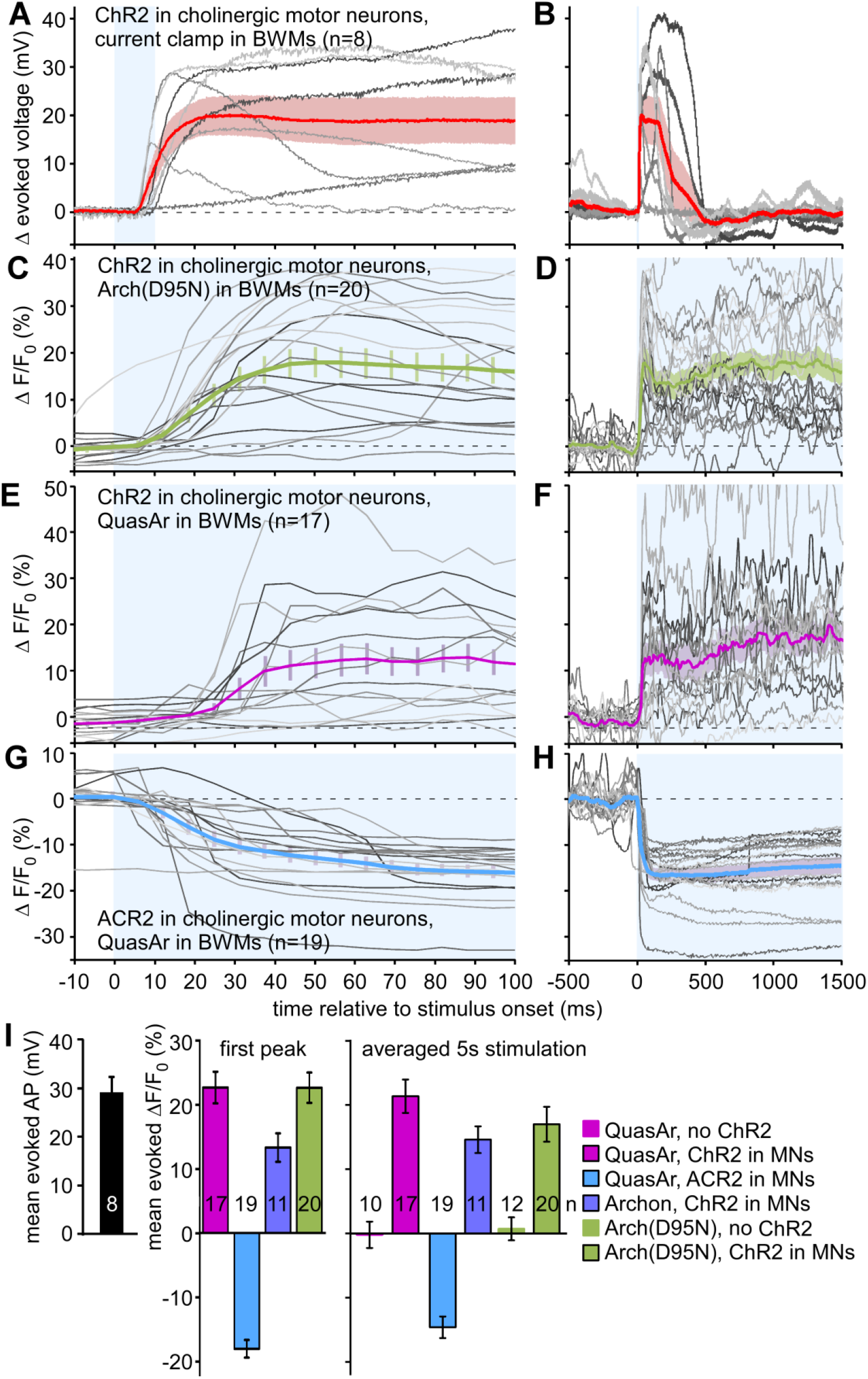
Calibration of electrical and rhodopsin voltage sensor signals in BWM cells, induced by optogenetic manipulation of cholinergic motor neurons: **A)** ChR2 mediated depolarization of cholinergic motor neurons (10 ms light pulse, indicated by blue shade) evokes action potentials (APs) in BWM cells, recorded under current clamp (n=8 animals, single records in grey). Mean±SEM voltage trace is shown in red and pink shade. **B)** As in (A), but extended time scale. **C, D)** Arch(D95N) fluorescence voltage signals recorded in response to 5 s light pulse for photodepolarization of cholinergic MNs by ChR2. Mean±SEM and single records from indicated number of animals. **E, F)** As in (C, D), but using QuasAr as voltage sensor in BWMs. **G, H)** Using QuasAr as sensor, and ACR2 anion channelrhodopsin for hyperpolarization of cholinergic MNs, exhibiting a drop in fluorescence (i.e. voltage), in response to the seizing of ACh signals from the MNs. **I)** Mean analysis of the data shown in (A-H), for the electrophysiologically measured AP amplitudes, as well as for the first fluorescence peak (assuming first AP) or during the entire 5 s light stimulus, for the indicated combinations of sensors, actuators, and controls. Frame rate in C-H was 158 fps, 2 ms exposure. Number of recorded animals is indicated. See also **Supp. Fig. 1** for a representation of all experiments performed, and for the entire duration of the recordings.

### Arch(D95N) shows robust fluorescence increases in pharyngeal muscle

Next, we assessed voltage-dependent changes in rhodopsin fluorescence in the pharynx. This muscular pump, used for feeding, exhibits APs and contractions. Arch(D95N), despite showing dim fluorescence, displayed strong fluorescence changes upon pharynx activity, on average ca. 122 % ΔF/F (measured across the entire organ; **Fig. 4A, D; Supp. Video 5**). The pharynx AP was previously reported to be between 80 and 115 mV ^29-31^ (**Fig. 4C**), on average 97 mV. Thus, in the pharynx, a 100 mV depolarization corresponds to ca. 126 % fluorescence increase, which is larger than reported for Arch(D95N) or QuasAr in mammalian cells and neurons^17,18^, and also largely exceeds the fluorescence increase observed in BWM cells upon excitation (**Fig. 3C, D, I**), for unknown reasons. The signal to noise ratio (SNR) was also very good (ca. 70; **Fig. 4D**). However, detecting Arch(D95N) fluorescence in BWM cells requires a strong excitation laser (we achieved up to 1800 mW/mm^2^ at 637 nm; in the pharynx, ca. 1/10^th^ of this intensity was sufficient) and an EMCCD camera, i.e. equipment not available in every lab. We thus also assessed the potential of DMAR-supplemented Arch(D95N) for voltage measurement. The raw fluorescence was largely increased (**Fig. 1D**) and could be visualized with a common sCMOS camera and excitation through an HBO lamp (30 mW/mm^2^, 620 nm). This excitation, however, did not allow recording APs well. Possibly, also Arch(D95N)-DMAR fluorescence must be highly excited to become voltage sensitive. Thus these measurements were performed with laser excitation (637 nm, ca. 180 mW/mm^2^, and an EMCCD camera). Arch(D95N)-DMAR exhibited activity-dependent fluorescence increases of about ca. 27 % (i.e. ca. 28 % / 100 mV), much smaller than for ATR-supplemented Arch(D95N), with a SNR of ca. 35 (**Fig. 4A, D; Supp. Video 6**). Signals were very uniform within a pump train in the same animal, and varied more between animals (**Supp. Fig. 2A, B**). Thus, while the signal change per mV is ca. 5-fold lower for DMAR than for ATR, the much increased detectability of the fluorescence makes this a valid alternative. We also tested analog VI in Arch wt voltage imaging. However, the fluorescence of this retinal analog did not show any voltage dependence (data not shown).

**Figure 4.**
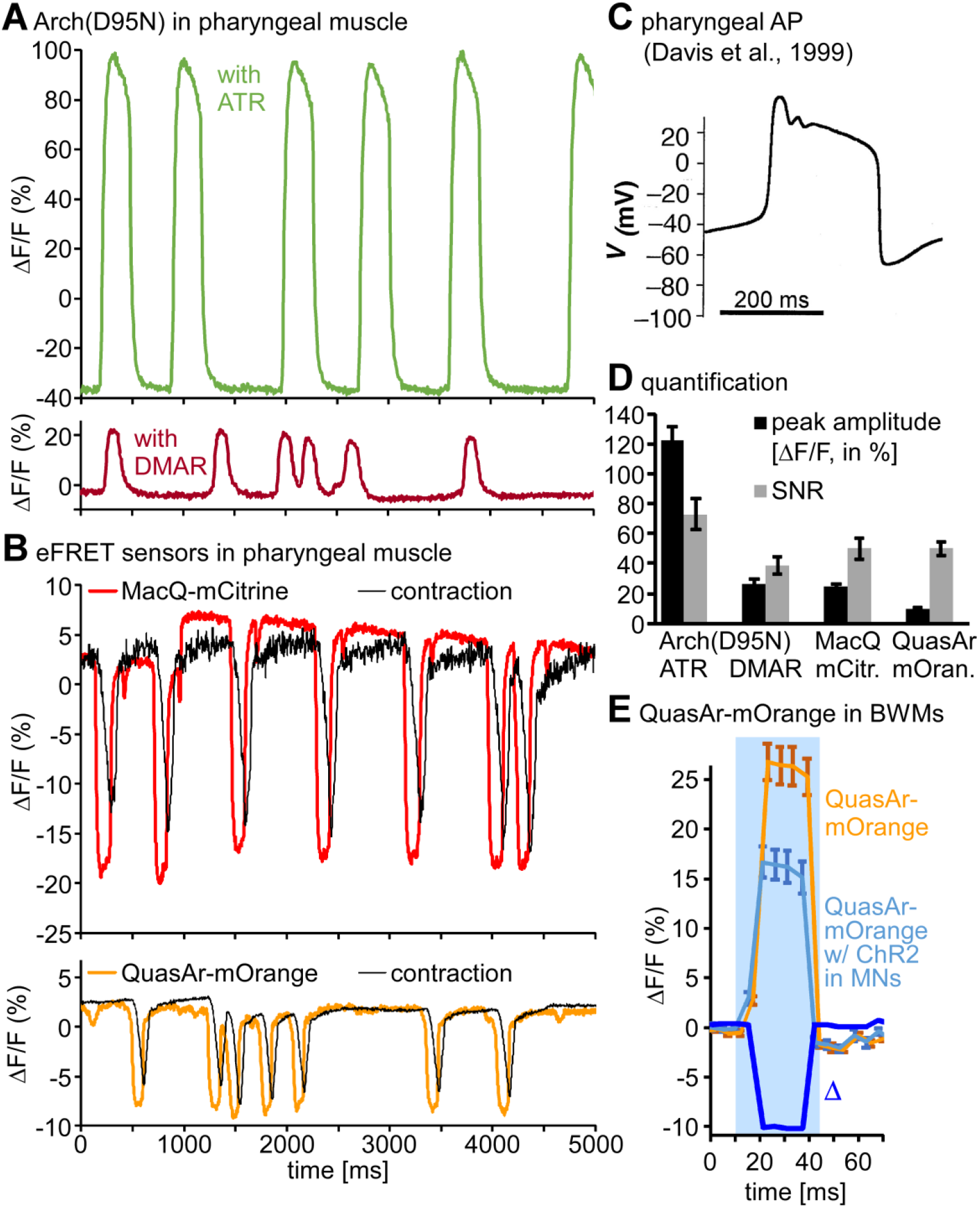
Arch(D95N) and eFRET voltage sensor signals quantified in pharyngeal muscles during spontaneous pumping: **A)** Arch(D95N) ΔF/F signals of pharyngeal muscles (entire organ) during pump trains, supplemented with ATR (upper panel) or DMAR (lower panel). See also **Supp. Videos 5 and 6**. **B)** Pharyngela APs evoked a drop in fluorescence when analyzed by eFRET sensors MacQ-mCitrine (upper panel, red; see also **Supp. Video 7**) and QuasAr-mOrange (lower panel, orange, see **Supp. Video 1**). Also shown is the corresponding contraction signal in the TB (black). **C)** Pharyngeal AP, measured by sharp electrode recording (reproduced, with permission, from Ref. 29). **D)** Group data for the sensors shown in (A, B), mean±SEM of the ΔF/F fluorescence peak amplitudes, as well as for the signal-to-noise ratio (SNR), defined by the ratio of peak amplitude divided by the standard deviation of the noise (fluorescence fluctuation between peaks). **E)** QuasAr-mOrange signal was measured in BWM cells in response to blue-light stimulation of ChR2 in cholinergic MN (bright blue graph) or without ChR2 expression (orange graph), before and during a blue light stimulus (blue shade). Note that both signals rise due to the additional excitation of mOrange by blue light. However, a difference graph (Δ, dark blue) shows a ca. 10% drop in fluorescence upon ChR2-mediated stimulation. n=7-8 animals with 3-15 APs each (or silent periods in between) were analyzed for (D), and n=12-13 animals were compared for the data in (E). Frame rate in (A, B, E): 189 fps, 1 ms exposure.

**Supplementary Video 5:** Arch(D95N)-ATR in pharynx, spontaneous pumping.

**Supplementary Video 6:** Arch(D95N)-DMAR in pharynx, spontaneous pumping.

**Supplementary Figure S2:**
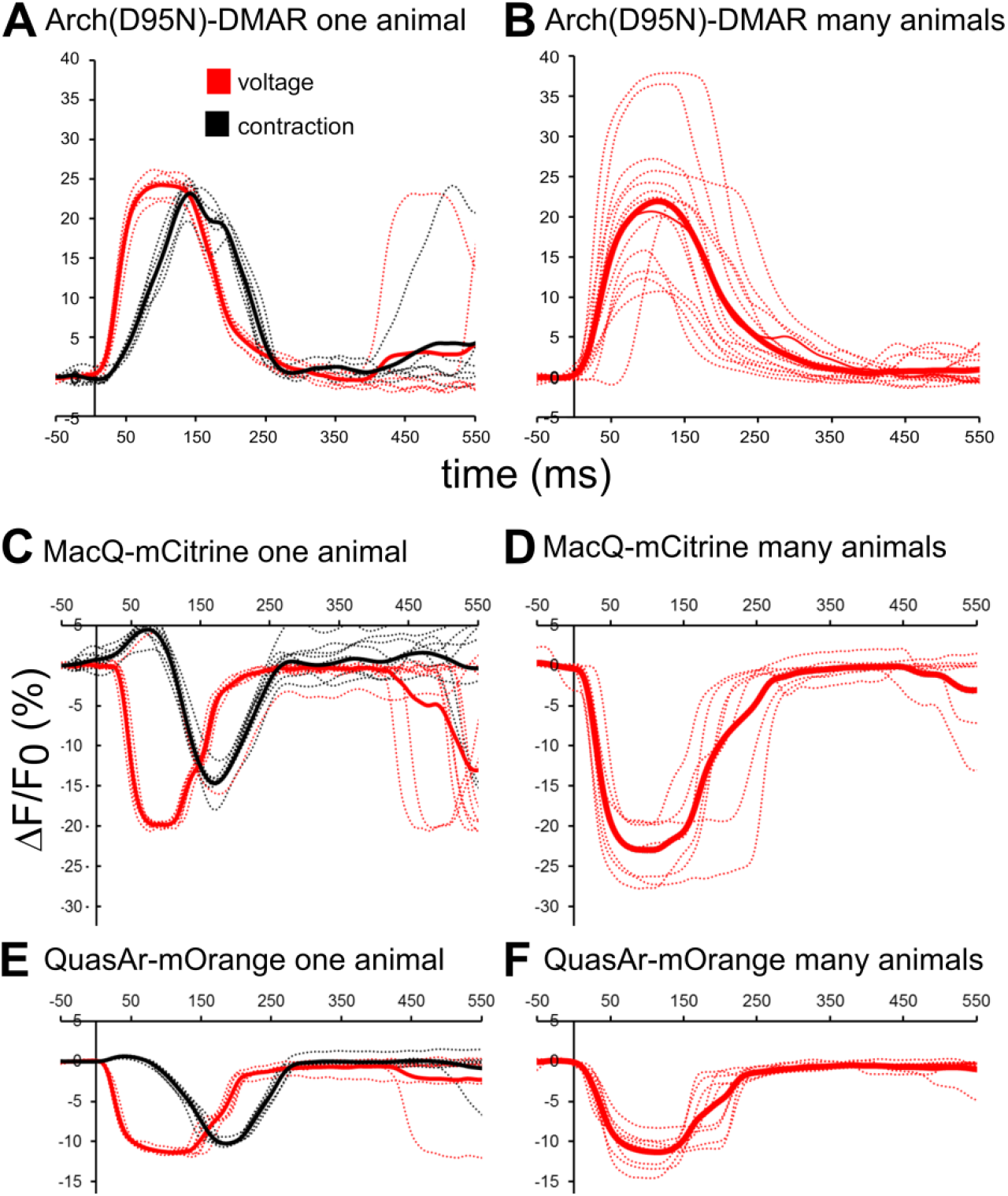
Comparison of voltage traces recorded using rhodopsin and eFRET voltage sensors in the pharynx during pump trains in single animals, and across animals. Related to Fig. 4. **A)** Arch(D95N)-DMAR, voltage and pump fluorescence signals of a single animal, aligned and averaged. Red: voltage trace; black: pump trace. **B)** As in (A), but mean pump events of 11 animals. **C, D)** Single animal events measured with MacQ-mCitrine in the pharynx (C) and mean events observed in 7 animals (D). **E, F)** As in (C, D), but using QuasAr-mOrange (n=7 animals in F).

### Electrochromic FRET sensors expressed in pharyngeal muscle report on voltage and enable concomitant analysis of muscle contraction

Last, we analyzed the eFRET sensors QuasAr-mOrange and MacQ-mCitrine in pharyngeal muscle cells during spontaneous pumping. For quantification we drew a region of interest (ROI) encircling the entire pharynx. A ROI outside the pharynx, but within the head region of the animal was analyzed for background signal. Fluorescence of both sensors could be readily visualized with standard epifluorescence microscope excitation intensities, as obtained from an HBO lamp or typical LEDs. MacQ-mCitrine and QuasAr-mOrange showed ca. −25 and ca. −10 % reductions in the fluorescence upon pharyngeal pumping, respectively (**Fig. 4B; Supp. Fig. 2C-F; Supp. Videos 1 and 7**). SNR was comparable for both sensors (ca. 50; **Fig. 4D**). Thus, despite lower overall fluorescence (**Fig. 1D**), MacQ-mCitrine is the more sensitive voltage reporter. The signals for both sensors were quite uniform in the same animal. However, mean signal traces between animals showed a range of amplitudes (**Supp. Fig. 2C-D**). Based on the reported pharyngeal AP, the signals of MacQ-mCitrine and QuasAr-mOrange correspond to a change in fluorescence of ≈26 and ≈10.4 % ΔF/F per 100 mV. For QuasAr-mOrange expressed in BWMs, we could more directly ‘calibrate’ this by assaying QuasAr-mCitrine signals observed with ChR2 stimulation in MNs, and comparing this to similarly illuminated animals lacking ChR2 expression (**Fig. 4E**): Although the sensor excitation light leads to a minor stimulation of ChR2, a blue-light pulse caused different fluorescence increases in the two strains (ca. 10 % smaller for animals expressing ChR2). As evoked APs are 29 mV, this corresponds to ca. −34 % ΔF/F_0_ per 100 mV. Thus, QuasAr-mOrange seems to be more sensitive in BWMs than in the pharynx.

**Supplementary Video 7:** MacQ-mCitrine in pharynx, spontaneous pumping.

Analyzing a small ROI enclosing the pharyngeal TB grinder region allowed deriving the opening of the pharynx from the fluorescence signal (**Figs. 2A, B; 4B**). Thus, we could correlate depolarization and the contraction of the muscle. We wrote scripts to systematically analyze the measured fluorescence traces. These involved automatic detection of relevant events, aligning, synchronizing, and averaging them. Then, parameters like amplitude, area under the peak, AP and pump duration, both defined as rise from and return to the baseline, or by using the full-width at half-maximum (FWHM), delay of pump vs. voltage, rise and drop of the voltage as τ_on_ and τ_on_ values, could be derived (**Fig. 5A**). This way, voltage and pump signals could be analyzed and compared for Arch(D95N) equipped with ATR and DMAR, MacQ-mCitrine, and QuasAr-mOrange (**Fig. 5B-H**; 8-15 animals with 2-10 APs, each).

**Figure 5.**
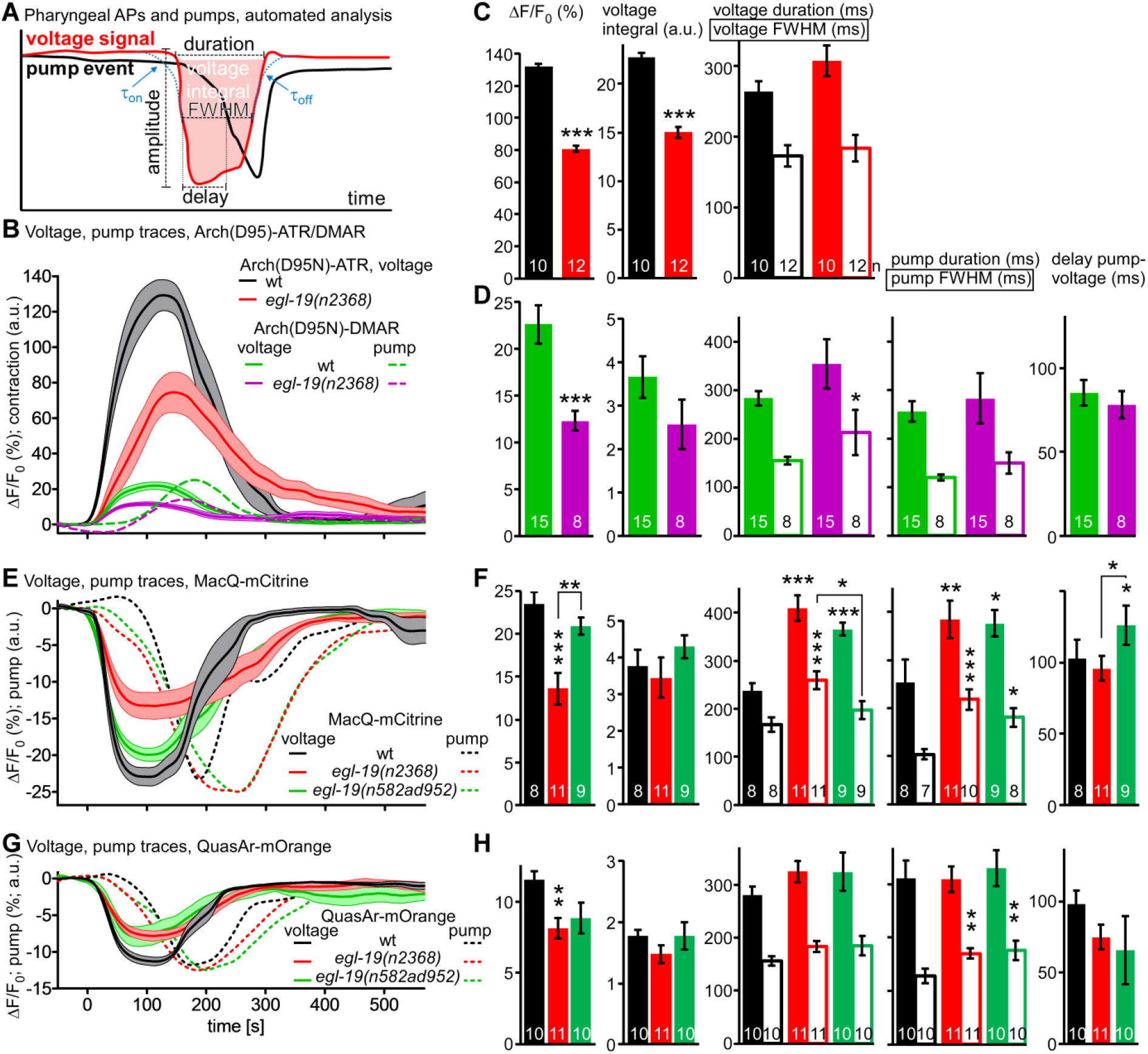
Pharyngeal AP and pump parameters quantified by automated analysis in wt and L-type VGCC g.o.f. mutants: **A)** Automated analysis of parameters of the pharyngeal AP and the corresponding pump contraction of the TB. Parameters determined after extraction and alignment of AP and pump events from fluorescence traces, using an automated analysis script (see methods for details). **B)** Mean±SEM data obtained after measurements using Arch(D95N) equipped with ATR and DMAR, in wt and in *egl-19(n2368)* mutants. **C, D)**. Parameters of APs and pumps deduced from the data shown in (B). **E, F)** Mean±SEM data and parameters deduced from MacQ-mCitrine recordings. **G, H)** Mean±SEM data and parameters deduced from QuasAr-mOrange recordings. Frame rate in (B, E, G): 189 fps, 1 ms exposure. Data in (C, D, F, H) were statistically analyzed by ANOVA, Bonferroni correction. ***P≤0.001, **P≤0.01, *P≤0.05.

Pumps could not be deduced for Arch(D95N)-ATR as the basal fluorescence was too dim to detect any decrease due to TB opening. A comparison between signals observed by the tested sensor-retinal combinations showed, as deduced from manual analysis, different amplitudes of the pharyngeal AP (132, 22.6, −23.5 and −11.6 % ΔF/F_0_ for Arch(D95N)-ATR, Arch(D95N)-DMAR, MacQ-mCitrine, and QuasAr-mOrange, respectively), while the duration of the AP was very similar (263.5, 283, 238 and 281 ms, respectively, no significant differences). For Arch(D95N)-DMAR, MacQ-mCitrine and QuasAr-mOrange, we could also compare other parameters among the sensors: The pump duration was 255, 264, and 304 ms, respectively (no significant difference); the delay from beginning of the voltage and pump FWHM was 85, 103 and 98 ms, respectively (no significant difference); voltage rise (ca. 15 ms) and decay (ca. 35 ms), as well as pump rise (erroneous for Arch(D95N), 23-39 ms for the eFRET sensors) and decay time constants (19-39 ms; **Supp. Fig. 3**) were not significantly different between the sensors. This demonstrates a tight coupling of voltage and contraction kinetics, observable with all of the sensors, and allows correlating voltage changes and physical contraction of this muscular structure. It should also allow to characterize aberrations of these times in mutants affecting pharyngeal muscle physiology, in ways not easily accessible by electrophysiology, as the movement of the muscles hinders precise intracellular recordings.

**Supplementary Figure S3:**
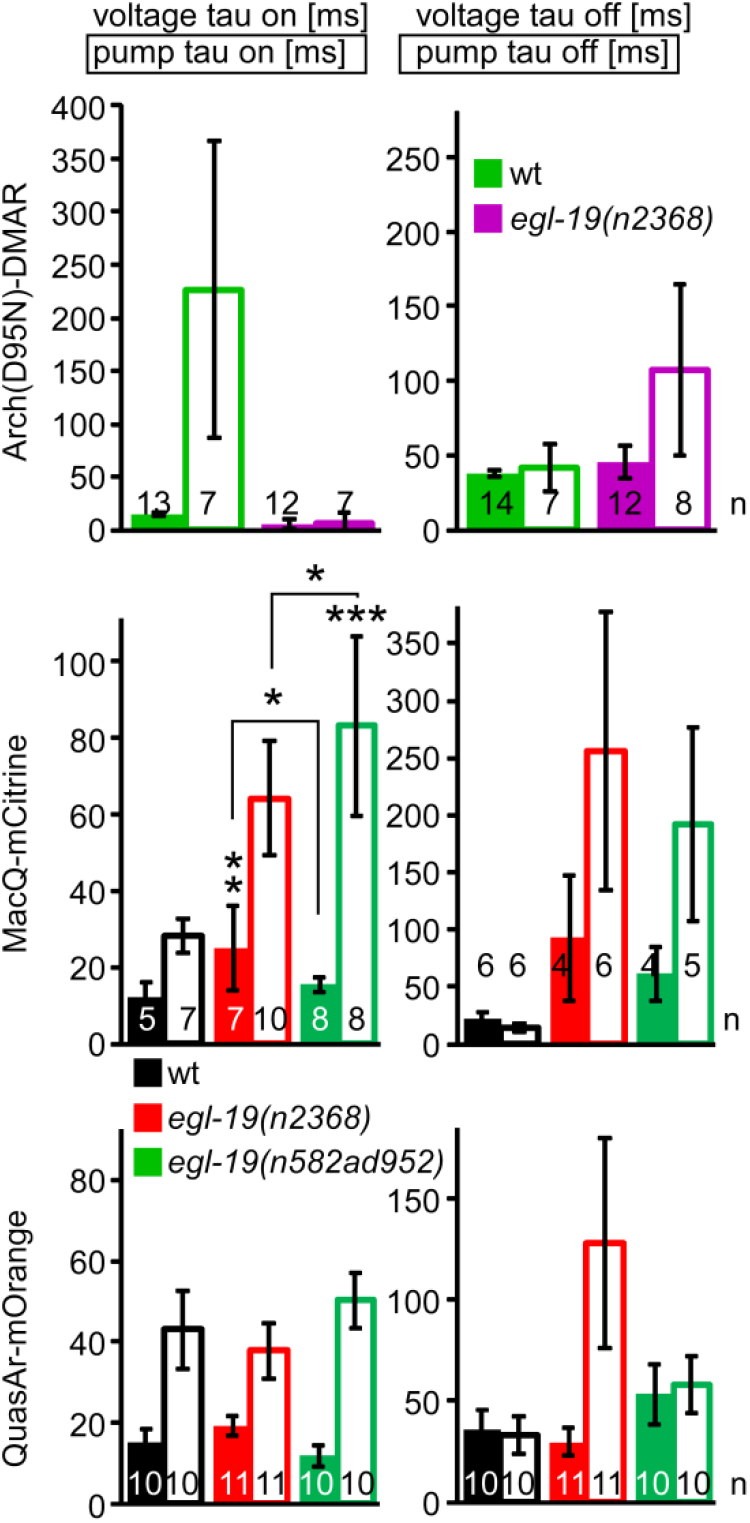
Voltage and pump signal rise and decay times, automatically determined for pharyngeal APs and pump events measured with rhodopsin and eFRET sensors. Related to Fig. 5. Parameters, genotypes and number of animals as indicated, measured for animals expressing Arch(D95N), equipped with DMAR (top), MacQ-mCitrine (middle) and QuasAr-mOrange (lower panels). Means±SEM, ANOVA with Bonferroni correction, ***P≤0.001, **P≤0.01, *P≤0.05.

### ‘Gain-of-function’ mutants in the L-type voltage gated Ca^2+^ channel EGL-19 exhibit reduced voltage increases, but prolonged pump duration

To see if the rhodopsin voltage sensors can be used to quantitatively compare mutants affecting APs, we focused on the L-type voltage gated Ca^2+^ channel (VGCC) EGL-19. This channel carries currents that shape the plateau phase of the pharyngeal AP^32,33^. Loss-of-function mutants are lethal, however, several ‘gain-of-function’ (g.o.f.) alleles have been isolated and characterized, e.g. by their effects on muscular tone (EGL-19 is also expressed in BWM cells). We analyzed two g.o.f. alleles, *n2368* and *n582ad952*. These affect amino acids close to the pore domain of the 2^nd^ module of the VGCC (*n2368* → G365R), or a double mutation in the pore of the 1^st^ and the voltage sensor of the 3^rd^ module (*n582ad952* → S372L, R899H). We compared 8-15 animals per genotype, averaging 2-10 APs per animal. Imaging pharynx pumping with either Arch(D95N)-ATR or -DMAR or the two eFRET voltage sensors showed a significantly reduced voltage amplitude signal for *n2368* (to 52-65 % of the wt; **Fig. 5B-H**), and a reduced signal (not significant) in the *n582ad952* allele. For the MacQ-mCitrine sensor, both alleles showed a significantly prolonged AP and pump duration (**Fig. 5E, F**; pump duration was also increased when measured by QuasAr-mOrange, **Fig. 5G, H**). Thus, the term ‘g.o.f.’ is mostly valid by the consequence of these alleles on BWM contraction properties (i.e. prolonged pump), but not for the AP (which in BWMs is likewise reduced in amplitude, though also prolonged^24,25^). Therefore, these alleles are hypomorphic rather than g.o.f., emphasizing that voltage imaging is a valuable extension of current methods for characterization of mutations in ion channels, affecting excitable cells. In particular, imaging is a more easily applied alternative to electrophysiology, which is experimentally demanding and not available in many labs.

It was previously shown that the dihydropyridine analog nemadipine A (NemA) is a potent inhibitor of EGL-19, or may accelerate its desensitization^34,35^. We analyzed whether nemadipine could influence the voltage-dependent fluorescence signals of the GEVIs tested in the pharynx. We compared the data from animals incubated in NemA to animals incubated in just the DMSO-containing vehicle (**Supp. Fig. 4**). As expected, NemA decreased wild type (wt) voltage signals, and essentially restored the *egl-19* g.o.f. alleles to DMSO-control levels in wt. This was true for all sensors tested (Arch(D95N)-DMAR, QuasAr-mOrange, and MacQ-mCitrine), although to different degrees.

**Supplementary Figure S4:**
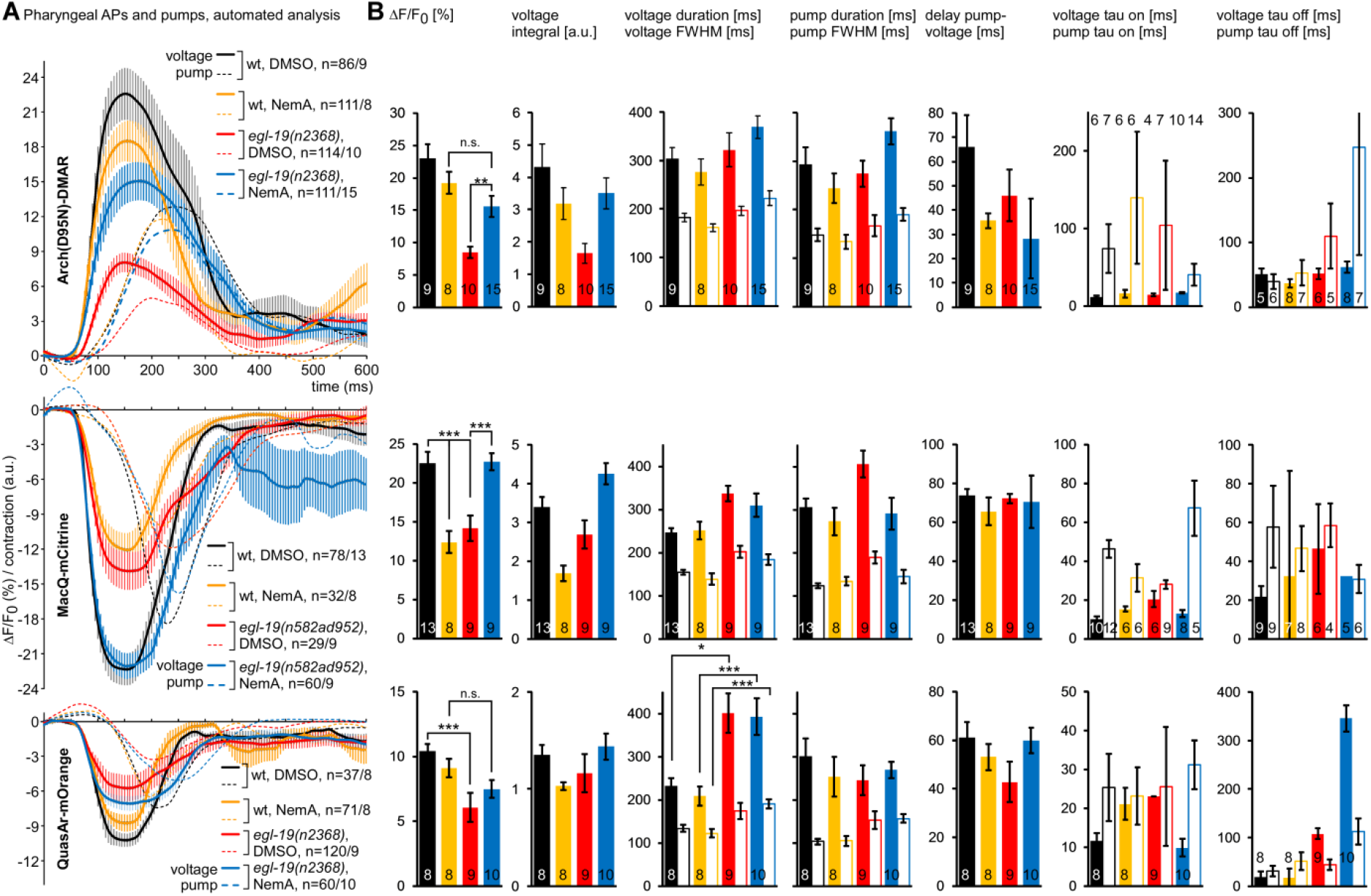
Comparison of voltage and pump parameters measured in mutants and in response to nemadipine-A (NemA) in the pharynx. **A)** Mean±SEM values for the indicated genotypes and drug or vehicle treatments, as well as different rhodopsin voltage sensors, as indicated. Also indicated is the number of animals analyzed, as well as the number of APs and pump events. **B)** Group data of automated analysis of parameters extracted from the data shown in (A), as in Fig. 5. Frame rate in (A): 189 fps, 1 ms exposure. Means±SEM, ANOVA with Bonferroni correction, ***P≤0.001, **P≤0.01, *P≤0.05.

### Imaging spatially compartmentalized voltage signals in excitable cell ensembles

Voltage recording can be done by electrophysiology with high accuracy. Yet, in electrically coupled cell systems, it is not possible to assess potentially compartmentalized electrical properties of the cell ensemble. This is expected to be the case for the pharynx, where anterior and posterior portions act differently, as was suggested for the different events visible in electropharyngeogram (EPG), i.e. the extracellular recording of pharyngeal electrical activities^36-38^. EPGs are current recordings with a complex, yet stereotypic structure (**Fig. 6C**, inset). They include currents corresponding to pharyngeal muscle depolarization (‘outward’ currents, as they correspond to current flowing from the extracellular space into the muscle) and repolarization events (i.e. ‘inward currents’). Also sometimes observed are currents corresponding to the activity of inhibitory neurons that terminate the pharyngeal AP; these show up as small inward currents, and may thus correspond to negative charge rushing into the muscle. Such events are not spatially resolvable by EPG recordings, and have not been addressed by sharp electrode recordings from different sections of the pharynx (only the TB has been recorded from)^29-32,39^. We thus explored if the imaging approach allows visualizing compartmentalized activity in the pharynx. To this end, we assessed a MacQ-mCitrine recording of a pump train with uniform single events (**Supp. Video 7**). To focus on changes in voltage, rather than absolute signals, we calculated difference videos, by deducing the value of every pixel from the value of the same pixel in the preceding video frame. After temporally aligning and averaging of twelve events, we could clearly observe depolarization and repolarization events (**Fig. 6A; Supp. Video 8**). The spatiotemporal differences in voltage change could be timed to single video frames and assessed in a kymograph along the longitudinal axis of the pharynx (**Fig. 6A, B**). The entire pharynx synchronously depolarized, while repolarization, following ca. 144 ms after the depolarization onset, occurred first in the anterior portion of the pharynx (corpus), and was followed 50 ms later by repolarization in the isthmus and TB. In the region connecting corpus and isthmus was a small section showing an additional depolarization between anterior and posterior repolarization phases (at 161 ms), and an additional repolarization phase following repolarization of the TB and isthmus. This region could match the connection between pm5 and pm4 muscle cells. Also, here are multiple inputs from pharyngeal neurons, thus possibly the minor signals occur from neuronal input to the pm5 muscle. It will be interesting to visualize this region in more detail, ideally with markers for pharyngeal neuron cell types.

**Figure 6.**
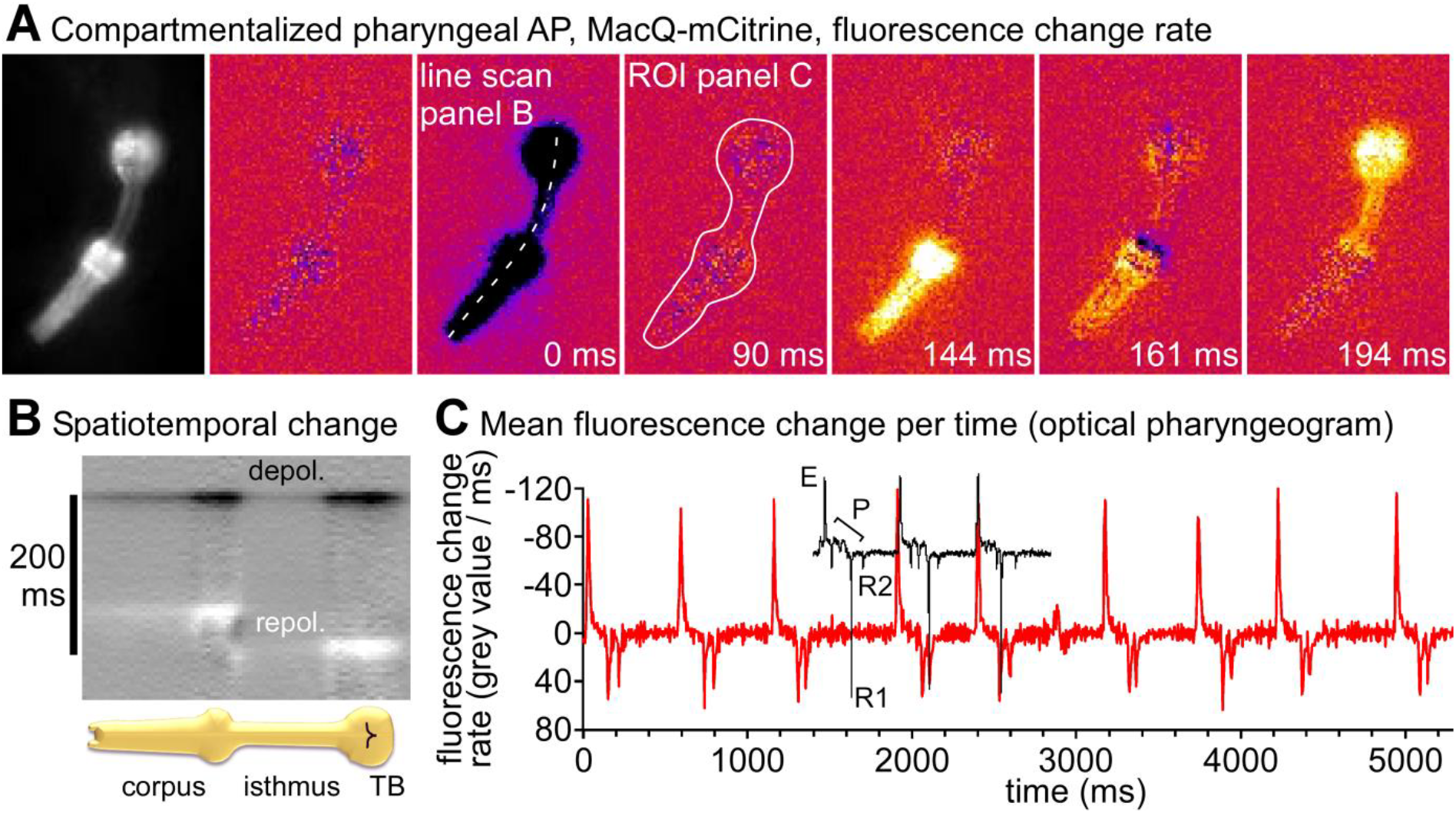
Pharyngeal AP repolarization occurs in a spatiotemporally compartmentalized fashion: **A)** A MacQ-mCitrine fluorescence video (191 fps, 1 ms exposure) of a pharyngeal pump train was analyzed by measuring fluorescence differences from frame to frame and averaging 12 events after alignment to the peak depolarization rate (0 ms, as indicated). Dark and bright colors represent high rates of depolarization and repolarization, respectively. See also **Supp. Video 8**. A line drawn along the axis of the pharynx allows generating a kymograph (**B**) for analysis of spatiotemporal development of the voltage along the pharynx, showing consecutive repolarization for corpus and terminal bulb. A ROI around the entire pharynx allows generating an ‘optical EPG’ (**C**) by plotting the inverted mean values. For illustration purposes, an example EPG trace is overlaid in black, showing (from left to right) 3 repeats of the typical sequence of EPG spikes: e/E1(not shown) and E/E2, inhibitory P spikes, as well as r and R (R1 and R2).

**Supplementary Video 8:** MacQ-mCitrine in pharynx, spontaneous pumping, difference video, 12 events aligned, averaged, 10x slowed.

The EPG measures currents from the extracellular medium into pharyngeal muscle. It thus represents changes in muscle polarization. We thus wondered if the mean voltage difference events, averaged across the whole pharynx, would correspond to the EPG. When we calculated the inverse mean signal change of the entire pharynx, it matched the time course of a typical EPG recording well (**Fig. 6C**, see inset for a typical EPG record). A single, major depolarization spike (corresponding to ‘E’ or ‘E2’) was followed by two repolarization spikes, corresponding to the anterior and posterior parts of the pharynx (‘R1’ and ‘R2’)^36,38-41^. Thus, the optical measurement matches the EPG measurement, but outperforms it with respect to spatial information.

## DISCUSSION

In this work, we surveyed a number of microbial rhodopsin-based optical voltage sensors in different muscular tissues of *C. elegans*. These include the rhodopsin variants Arch(D95N)(equipped with ATR and the retinal analog DMAR), QuasAr, and Archon, as well as the eFRET sensors MacQ-mCitrine and QuasAr-mOrange. We show that all of them can be used to detect action potentials with robust fluorescence changes of up to 126 % per 100 mV in the pharynx, and 78 % in body wall muscles. These values were larger than previously reported for cells and neurons of other organisms; it may have to do with the lipid composition of the *C. elegans* cell membranes, which might influence the properties of the Arch protein in a beneficial manner. Making these high ΔF/F_0_ values accessible, however, requires the use of a high-power excitation laser (up to 1800 mW/mm^2^ 637 nm light), that are achieved only in a small field of view, and an EMCCD camera, to enable recording at high frame rate despite the low absolute fluorescence. We note that the addition of ATR to the culture medium caused increased background fluorescence upon blue light stimulation, thus low concentrations of ATR are recommended. Lower excitation light intensities (180 mW/mm^2^) were possible by using DMAR as an alternative to ATR. This strongly boosted the absolute fluorescence levels, yet reduced the fluorescence change per voltage change by a factor of 5. Possibly, a similar branched photocycle as in Arch(D95N)-ATR^20^ is present with DMAR, requiring more than one photon to be absorbed. Nonetheless, it was feasible to use this sensor/retinal analog combination to robustly measure parameters of pharyngeal pumping that were comparable to those measured with the eFRET sensors, and to compare VGCC g.o.f. mutants to wt.

The MacQ-mCitrine eFRET sensor, despite 75 % lower absolute fluorescence intensities, showed ca. 3-fold higher ΔF/F_0_ values than QuasAr-mOrange. Parameters of pharyngeal APs and pumping were similar for all sensors, emphasizing that they did not alter the properties of the pharynx relative to each other. Electrophysiological measurements are demanding and do not allow to analyze muscular movement and contraction parameters concomitantly, while the imaging approach shown here does. Since the parameters we compared between wt and *egl-19* g.o.f. alleles did not differ between the three sensors, it can be expected that also other mutations affecting pharyngeal physiology and pumping will be accessible using the methods described here. Furthermore, we showed that nemadipine A, a VGCC blocker, similarly affected - actually restored - pharyngeal AP characteristics in *egl-19* g.o.f. mutants, as shown earlier by electrophysiology^34^. The signals we determined for pharyngeal voltage fluorescence fluctuations were quite comparable in pump trains of single animals, but differed across comparisons between different animals. This may be due to the differences in immobilization of each individual, and this may be improved by more reproducible immobilization conditions as in a microfluidic device.

The imaging approach also allows following fluorescence fluctuations in a spatiotemporally defined manner across an ensemble of electrically coupled cells. This confirmed earlier assumptions made from EPG recordings, i.e. that the terminal bulb repolarization is delayed relative to the corpus repolarization. Yet, this was never simultaneously measured by sharp electrodes in the two parts of the pharynx, emphasizing the new possibilities provided by the optical measurement. Here, we observed not only the functional electrical compartmentalization of the two major parts of the pharynx, but also observed smaller sub-compartments at the interface of these two parts of the pharynx with potentially distinct electrical activities during distinct phases of the pharyngeal excitation-repolarization cycle. This could be further assessed, e.g. in gap junction mutants or cell-specific knock-down animals. Evaluating voltage change rates across the pharynx allowed to generate an ‘optical EPG’ that recapitulated most of the characteristic features of the classical EPG.

The combination of voltage imaging in the infrared spectral range and blue-light stimulation of ChR2 in excitable cells enables all-optical electrophysiology approaches for *C. elegans*, and will greatly facilitate such measurements without the need for complicated electrophysiological approaches^42^. This should enable to analyze mutants affecting neuromuscular junction function that are too small to allow dissection needed for patch-clamp electrophysiology. Additionally, the fact that voltage imaging is performed in intact animals is a distinct advantage over measurements in dissected animals, where the composition of the extracellular and pipette solutions may alter the endogenous physiology. Future work will use the methods established here for *C. elegans* neurons, which are even more complicated to address by electrophysiology. In sum, we compared a range of useful tools for all-optical interrogation of muscular tissues and ensembles, in response to intrinsic activities, as well as optogenetically evoked neurotransmission in the nematode.

## METHODS

### *C. elegans* culture and transgenic animals

Worms were cultivated on nematode growth medium (NGM) plates, seeded with *E. coli* OP50-1 strain, in 6 cm petri dishes, as described previously^42^. Generation of transgenic animals was performed by microinjection of plasmid DNA, at varying concentrations^43^. For transgenic strains ZX1917, ZX1918 and ZX1958, 3 ng/µL of the respective voltage sensor plasmid DNA was injected into the gonads of mother animals. For strains ZX1920, ZX1953, ZX2476, ZX2479 and ZX2483, 10 ng/µL of the respective voltage sensor plasmid DNA were used in combination with either 1.5 ng/µL of the co-expression marker p*myo-2*::CFP or 30 ng/µL of p*elt-2*::GFP or p*myo-3*::CFP. Additionally, for strain ZX2483 50 ng/µL of p*unc-17*::ACR2::eYFP were injected.

#### *Transgenic* C. elegans *strains*

The following strains were used or generated: **DA95:** *egl-19(n582ad952)*, **MT6129**: *egl-19(n2368)*, **ZX1907**: *zxEx922[pmyo2::Arch::2xMyc; pmyo-3::CFP]*, **ZX1917**: *zxEx942[pmyo-2::QuasAr::mOrange; pmyo-3::CFP]*, **ZX1918**: *zxEx943[pmyo-2::MacQ::mCitrine; pmyo-3::mCherry]*, **ZX1920:** *zxIs121[pmyo-3::QuasAr::mOrange; pmyo-3::CFP]*, **ZX1951**: *egl-19(n2368); zxEx942[pmyo-2::QuasAr::mOrange; pmyo-3::CFP]*, **ZX1952**: *egl-19(n2368); zxEx942[pmyo-2::MacQ-mCitrine; pmyo-3::mCherry]*, **ZX1953:** *zxIs120[pmyo-3::ArchD95N::2xmyc-tag; pmyo-3::CFP]*, **ZX1954:** *zxIs6; zxIs120[pmyo-3::ArchD95N::2xmyc-tag; pmyo-3::CFP]*, **ZX1955**: *egl-19(n582ad952); zxEx942[pmyo-2::QuasAr::mOrange; pmyo-3::CFP]*, **ZX1956**: *egl-19(n582ad952); zxEx942[pmyo-2::MacQ-mCitrine; pmyo-3::mCherry]*, **ZX1958**: *zxEx944[pmyo-2::ArchD95N::2xmyc-tag; pmyo3::CFP]*, **ZX1959**: *egl-19(n582ad952); zxEx944[pmyo-2::ArchD95N; pmyo-3::CFP]*, **ZX1960**: *zxIs5[punc-17::ChR2(H134R); lin-15+]; zxIs121[pmyo-3::QuasAr::mOrange; pmyo-3::CFP]*, **ZX2476**: *zxEx1139[pmyo-3::QuasAr; pmyo-2::CFP]*, **ZX2479**: *zxIs5[punc-17::ChR2(H134R); lin-15+]; zxEx1142[pmyo-3::Archon::GFP; pmyo-2::CFP]*, **ZX2482**: *zxIs5[punc-17::ChR2(H134R); lin-15+]; zxEx1145[pmyo-3::QuasAr; pmyo-2::CFP]*, **ZX2483**: *zxEx1146[punc-17::ACR2::eYFP; pmyo-3::QuasAr; pelt-2::GFP]*.

### Molecular biology

Plasmid pAB4 (*punc-17::ACR2::eYFP*) was already described previously (Ref. 27).

Plasmid pAB16 (p*myo-3*::QuasAr) was generated via Gibson Assembly based on p*myo-3*::QuasAr::mOrange and pPD96.52 (p*myo-3*, Fire Lab Vector Kit, Addgene plasmid #1608), using restriction enzymes *Bam*HI and *Xba*I and primers QUASAR_fwd (5’-cccacgaccactagatccatATGGTAAGTATCGCTCTGCAG-3’) and QUASAR_rev (5’-gtcctttggccaatcccgggCTCGGTGTCGCCCAGAATAG-3’). Plasmid pAB19 (p*myo-3*::Archon::GFP) was generated via subcloning from *rig-3*::wArchon1::GFP (a gift from Steven Flavell) into pPD96.52, using restriction enzymes *Kpn*I und *Sac*I.

Plasmids pNH13 (p*myo-2*::QuasAr::mOrange) and pNH12 (p*myo-2*::MacQ::mCitrine) were generated by subcloning of plasmids #59173 and #48762 (Addgene) into pPD132.102 (*pmyo-2*, #1662, Addgene) via restriction with *Bam*HI and *Eco*RI. A site-directed mutagenesis of plasmid pNH10 (p*myo-2*::Arch::2xMycTag) resulted in plasmid pNH11 (p*myo-2*::Arch(D95N)::2xMycTag), using primers oNH7 (5’TATGCCAGGTACGCCAACTGGCTGTTTACCACC3’) and oNH8 (5’-GGTGGTAAACAGCCAGTCGGCGTACCTGGCATA-3’).

### Voltage imaging of immobilized animals

One day prior to the experiments, transgenic animals at L4 stage were transferred onto NGM plates seeded with OP50-1 lawns, grown from bacterial suspensions supplemented with ATR (Sigma-Aldrich, USA) or the respective ATR-analogues DMAR or VI (final ATR concentration: 0.01 mM for experiments with ChR2 or ACR2 blue-light stimulation, 1 mM for imaging experiments or Arch(D95N) in the pharynx, and 0.1 mM for imaging eFRET sensors); note that for analog VI we used Arch wt, an HBO lamp (30 mW/mm^2^ excitation at 620 nm; analog DMAR could be visualized well with HBO illumination, however, APs were observed clearly only with 637 nm laser excitation, 180 mW/mm^2^). For imaging, worms were placed onto 10 % agarose pads (dissolved in M9 buffer) on microscope slides and immobilized with polystyrene beads (0.1 µm diameter, at 2.5% w/v, Sigma-Aldrich). Imaging was performed on an inverted microscope (Zeiss Axio Observer Z1), equipped with an 40x oil immersion objective (Zeiss EC Plan-NEOFLUAR 40x/ N.A. 1.3, Oil DIC ∞ / 0.17), a 470 nm LED light source (KSL 70, Rapp OptoElectronic, Hamburg, Germany) or monochromator (PolychromeV, Till Photonics) for photostimulation, a 637 nm laser (OBIS FP 637LX, Coherent), a galilean beam expander (BE02-05-A, Thorlabs) for excitation of voltage sensors, and an EMCCD Camera (Evolve 512 Delta, Photometrics). Voltage sensor fluorescence was imaged at 637 nm excitation (1800 mW/mm^2^ for direct imaging of QuasAr and Archon in BWMs and ca. 180 mW/mm^2^ for pharynx measurements and direct imaging of Arch(D95N)-DMAR in BWMs) and 700 or 747 nm emission (700/75 ET Bandpass filter, integrated in Cy5 filter cube, AHF Analysentechnik) for ATR, and at 780 nm (780 LP ET Longpass Filter) for DMAR. eFRET sensors were imaged with 545/30 or 472/30 nm excitation (HBO100 100W mercury lamp, Zeiss, at 30 mW/mm^2^ for 545/30 nm and 10 mW/mm^2^ for 472/30 nm) and 610/75 or 520/35 nm emission (472/30 ET Bandpass, 610/75 ET Bandpass and 520/35 ET Bandpass Filters, AHF Analysentechnik), respectively. Stimulation of optogenetic actuators (ChR2 or ACR2) was performed with 300 µW/mm^2^ at 470 nm. Acquisition times, frame rates and gain used for imaging rhodopsin voltage sensors were 1 or 2 ms, 158 or 190 fps and gain 1 (for DMAR) or gain 10 or 50 (for ATR). For eFRET sensors, it was 1 ms exposure, ca. 190 fps and gain 10, if not otherwise stated.

To induce pharyngeal pumping under imaging conditions, the animals were incubated in 3 µl serotonin (20 mM, in M9 buffer) for 3 minutes immediately prior to the experiments. For drug testing, animals were exposed to either 30 nM nemadipine A, dissolved in 0.2 % DMSO in M9 or 0.2 % DMSO (vehicle control, in M9 buffer) respectively.

### Image acquisition, processing and automated data analysis of pharyngeal voltage and pump events

Video acquisition (with frame rates of 160 or ca. 190 fps, at 1 ms exposure time) was performed by using the *Photometrics PVCAM Device Adapter for µManager*^44^. *For BWM experiments, raw image sequences were analyzed via ROI (region of interest) selection of signal and background and multi-measure* function in ImageJ (National Institutes of Health, USA; https://imagej.nih.gov/ij/index.html). Changes in fluorescence (ΔF/F_0_, or ΔF/F_mean_, where F_mean_ (or F, for simplicity) is the mean fluorescence (F_mean_) of the ROI, calculated for the entire imaging period) were calculated as follows:

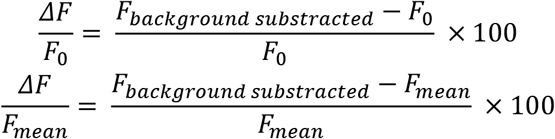

Contrast was calculated as the ratio between mean fluorescence of a ROI across the pharyngeal or BWM tissue expressing the voltage sensor, and the mean fluorescence of a ROI within the animal, but chosen in tissue not expressing sensors and showing no autofluorescence.

Signal to noise ratio (SNR) was calculated as the ratio of the mean fluorescence of all peaks (= APs) observed during a pump train, and the standard deviation of the fluorescence in the ROI during resting phases.

### Automated analysis of voltage and pump signals in the pharynx

To register the voltage signals in the pharyngeal muscles, a ROI was defined around the whole pharynx (Fig. 2A), whereas a smaller circular ROI was positioned over the terminal bulb lumen with the grinder, to track grinder movements and contractions with the accompanying lumen opening (observed as a lowering of the fluorescence in this ROI, as fluorescing tissue moves radially away during TB contraction). For background correction, another circular ROI was defined in an arbitrary dark region outside of the worm, or, for contrast measurements, ‘inside’ the worm, but covering tissue that is not expressing the respective voltage sensor (and avoiding the autofluorescing gut). ΔF/F was calculated in Excel (Microsoft). A custom workflow in KNIME 3.6.2 (KNIME AG, Zurich, Switzerland^45^) was used to synchronize pump and voltage events across animals by calling an R (R 3.5.1, The R Foundation for Statistical Computing) script to process each of the Excel tables. The R script proceeds to fit a spline curve to the input data with as many anchor points as data points present. The fit curve is sampled at constant 200 data points per second to account for the variable camera timing. Pumps are found as local minima with a centered time window of 625 ms. A manual input of indices can be used to correct this peak registration. The signals that passed manual control are synchronized first to the pump minimum. The voltage signal is then analyzed for the peak onset by searching for the minimum of the scaled difference with a lag of 50 ms of a centered moving average with window size of 105 ms. The events are subsequently synchronized to the onset of the voltage peak and grouped per animal, or analyzed individually, where appropriate. The signals are further analyzed to extract start and end of peak by searching for the first and last data point to cross a line at 10 % of the distance from the baseline to the peak minimum, respectively. Analogously, the full width at half maximum (FWHM) is calculated by searching for the two data points that cross the 50 % line between baseline and peak minimum. Duration is defined as the time difference between start and end of a peak. The kinetic parameters τ_ON_ and τ_OFF_ are modelled each with a mono-exponential curve fit accounting only for the time frames from (−50 ms, FWHM_start_) and (FWHM_end_, 305 ms), respectively. The area (voltage integral) is calculated as the integral from start to end of the peak from the baseline. The amplitude is calculated as the peak minimum to the maximum value at negative time after synchronization in the depicted time frame. The delay is reported as the time difference between the FWHM_start_ of the voltage and FWHM_start_ of the pump peaks.

## Electrophysiology

Recordings were conducted from dissected BWM cells on the ventral side of the body anterior to the vulva as described previously^42^. Animals were immobilized with Histoacryl glue (B. Braun Surgical, Spain) and a lateral incision was made to access neuromuscular junctions along the ventral nerve cord. The basement membrane overlying muscles was enzymatically removed by incubation in 0.5 mg/ml collagenase for 10 s (C5138, Sigma-Aldrich, Germany). BWMs were patch-clamped in whole-cell mode at 22°C using an EPC10 amplifier with head stage connected to a standard HEKA pipette holder for fire-polished borosilicate pipettes (1B100F-4, WPI, USA) of 4-7 MΩ resistance. The extracellular (bath) solution contained: NaCl 150 mM; KCl 5 mM; CaCl_2_ 5 mM; MgCl_2_ 1 mM; glucose 10 mM; sucrose 5 mM; HEPES 15 mM, pH7.3 with NaOH, ≈330 mOsm. The pipette solution contained: K-gluconate 115 mM; KCl 25 mM; CaCl_2_ 0.1 mM; MgCl_2_ 5 mM; BAPTA 1 mM; Hepes 10 mM; Na_2_ATP 5 mM; Na_2_GTP 0.5 mM; cAMP 0.5 mM; cGMP 0.5 mM, pH7.2 with KOH, ≈320mOsm. Recordings were conducted in current clamp mode using Patchmaster software (HEKA, Germany), as described previously^27^. Light activation was performed using a LED lamp (KSL-70, Rapp OptoElectronic, Hamburg, Germany; 470 nm, 1 mW/mm^2^) and controlled by the EPC10 amplifier. Data were analyzed by Patchmaster software.

## Software

Knime 3.6.2 (KNIME AG, Zurich, CH), R 3.5.1 (The R Foundation for Statistical Computing)

## DATA AVAILABILITY

The original data, as videos or fluorescence grey values of the relevant ROIs, as well as the KNIME and R scripts used for the data analysis, will be made available through a publicly accessible server.

## AUTHOR CONTRIBUTIONS

Conceived experiments and analyzed data: NAH, ACFB, RS, WSC, JL, AG; planned and conceptualized research: NAH, ACFB, RS, WSC, JL, AG; designed and programmed software: WSC; wrote the paper, with the help of the other authors: AG; acquired funding: AG.

## ACKNOWLEDGEMENTS

We thank Steven Flavell and Adam Cohen for reagents, and Alexander Hirschhäuser, Heike Fettermann, Mona Hoeret, Regina Wagner and Franziska Baumbach for expert technical assistance. We are grateful to Christina Schüler for providing an original EPG record. DMAR and retinal Analog VI were gifts by Lars Kattner (Endotherm company). Some strains were provided by the *Caenorhabditis* Genetics Center, which is funded by the NIH - Office of Research Infrastructure Programs (grant P40 OD010440). This work was funded by Goethe University and the Deutsche Forschungsgemeinschaft (DFG), grants SFB807-P11 and EXC115 (Cluster of Excellence Frankfurt - Macromolecular Complexes) to A.G., and by an IMPReS PhD stipend to A.C.F.B.

